# Spacer2PAM: A computational framework for identification of functional PAM sequences for endogenous CRISPR systems

**DOI:** 10.1101/2021.08.20.457124

**Authors:** Grant A. Rybnicky, Nicholas A. Fackler, Ashty S. Karim, Michael Köpke, Michael C. Jewett

## Abstract

RNA-guided nucleases from clustered regularly interspaced short palindromic repeats (CRISPR) systems expand opportunities for precise, targeted genome modification. Endogenous CRISPR systems in many bacteria and archaea are particularly attractive to circumvent expression, functionality, and unintended activity hurdles posed by heterologous CRISPR effectors. However, each CRISPR system recognizes a unique set of PAM sequences, which requires extensive screening of randomized DNA libraries. This challenge makes it difficult to develop endogenous CRISPR systems, especially in organisms that are slow-growing or have transformation idiosyncrasies. To address this limitation, we present Spacer2PAM, an easy-to-use, easy-to-interpret R package built to identify potential PAM sequences for any CRISPR system given its corresponding CRISPR array as input. Spacer2PAM can be used in “Quick” mode to generate a single PAM prediction that is likely to be functional or in “Comprehensive” mode to inform targeted, unpooled PAM libraries small enough to screen in difficult to transform organisms. We demonstrate Spacer2PAM by predicting PAM sequences for industrially relevant organisms and experimentally identifying seven PAM sequences that mediate interference from the Spacer2PAM-predicted PAM library for the type I-B CRISPR system from *Clostridium autoethanogenum*. We anticipate that Spacer2PAM will facilitate the use of endogenous CRISPR systems for industrial biotechnology and synthetic biology.

## Introduction

Clustered regularly interspaced short palindromic repeats (CRISPR) system-derived, RNA-guided nucleases have enabled an abundance of technologies(1–3), including gene editing. While CRISPR gene editing within eukaryotes using heterologous components, like *Streptococcus pyogenes* Cas9, proves effective across eukaryotic phylogenetic space(4), success of those same components remains unpredictable across prokaryotes(5–8). In fact, use of heterologous CRISPR effectors in prokaryotes poses three main hurdles. First, transformation and expression of functional effector proteins is difficult in many non-model prokaryotes. Many common CRISPR effectors are large in size requiring over 3 kb of DNA sequence to encode the expression construct which can further reduce already low transformation efficiencies(9). Thus, using these effectors decreases the chance of successful transformation before the editing event even takes place. Second, the functionality of heterologous effector complexes is not guaranteed in the target organism’s cytosolic conditions. Enzymes are environmentally sensitive and demonstrate optimal activity within narrow physiological conditions. For example, the warm environment required by thermophiles can lead to inactivity of *S. pyogenes* Cas9(10). Third, CRISPR effectors have the potential to demonstrate off target activities or unexplained toxicities. Heterologous CRISPR effectors can possess additional activities that can interfere with gene editing or viability in prokaryotes(5–8, 11, 12) because CRISPR effectors are often sourced from other prokaryotic systems. Taken together, these hurdles make difficult the adoption of CRISPR gene editing in the growing listing of model and non-model prokaryotes relevant to industrial biotechnology and synthetic biology.

Endogenous CRISPR systems prevalent throughout bacteria and archaea(13) inherently avoid many of the barriers to using heterologous CRISPR effectors. Native systems are encoded within the genome and are often constitutively expressed(14, 15), adapted to function within their genome’s cytosolic environment(16), and have evolved to interact with their genome’s proteome without significant negative effects. In essence, using endogenous CRISPR systems presents unique opportunities for genome editing(14–18) and targeted antimicrobial applications(19–21) that otherwise would be inaccessible with current heterologous CRISPR effectors. However, identification of a functional protospacer adjacent motif (PAM) required for types I, II, and V CRISPR systems to target DNA(22) remains challenging when using endogenous CRISPR effectors. CRISPR effector complexes recognize a unique PAM or set of PAM sequences that is not easily gleaned from readily available information such as host organism or comparative genomics. Functional PAM identification thus requires empirical determination for each endogenous CRISPR system.

Current methods of PAM determination are often difficult to apply to CRISPR systems in prokaryotes without robust genetic tools. The primary experimental method used to determine functional PAM sequences is the screening of a randomized, pooled PAM library in the organism encoding the CRISPR system(16). The library is sequenced before and after selection by the CRISPR system and the change in frequency of each PAM is calculated. Decreases in PAM frequencies are associated with successful targeting by the CRISPR system. Similarly, cell-free(23) and *in vivo*(24) heterologous expression of CRISPR effectors have been used to reconstitute CRISPR effectors and screen their PAM specificity. Alternatively, researchers with limited resources or organisms that do not transform well enough to screen a randomized, pooled PAM library screen an unpooled PAM library(17). The unpooled nature of the library circumvents the need for large numbers of transformants but limits the throughput of PAM sequences that can be screened.

Computational methods can bypass the need for efficient DNA transformation to identify PAM sequences. Rather than observe the interference activity of a CRISPR system biochemically, computational methods can back trace the spacer adaptation process bioinformatically. Where a CRISPR system naturally samples invading nucleic acids for the presence of a PAM before integrating protospacer into the CRISPR array(25), nucleotide alignment can be used to identify the origin of CRISPR array spacers and the sequence adjacent to the alignment can be queried for the identity of potential PAMs. By doing this process across all the spacers encoded by a CRISPR system’s arrays, the potential PAM sequences can be used to predict PAM preferences of that CRISPR system. Attempts at this process have been developed(17, 26, 27) but are often limited in their ability to identify functional PAMs, difficult to interpret into actionable experiments, or incomplete and require the use of multiple tools in a non-consolidated pipeline.

In this work, we develop, optimize, and apply Spacer2PAM, an R package built to identify functional PAM sequences for any CRISPR system given its corresponding CRISPR array as input. This tool improves upon previous computational methods by implementing filter criteria to down select the number of sequence alignments, generating a more biologically relevant set of candidate PAM sequences and increasing the frequency of functional PAM predictions. We validate Spacer2PAM with 10 well-characterized CRISPR systems and optimize Spacer2PAM to output an experimentally actionable consensus PAM sequence, a score for the PAM prediction, and an optional sequence logo representing the sequences used to build the consensus. We then apply Spacer2PAM to predict PAM sequences for type I-B CRISPR systems from 11 organisms with uncommon carbon metabolism. Further, we use these predictions to determine and experimentally validate functional PAMs for the *Clostridium autoethanogenum* type I-B CRISPR system. Spacer2PAM offers an easy-to-use computational tool for PAM prediction that we anticipate will facilitate research into novel CRISPR systems and spur new synthetic biology applications.

## Materials and Methods

### Prediction of PAM Sequences

All CRISPR arrays were retrieved from CRISPRCasdb, part of CRISPR-Cas++, which can be found at https://crisprcas.i2bc.paris-saclay.fr/ (28). Alignment of CRISPR spacers to genomes was done via the NCBI BLAST web interface(29) using the BLASTn algorithm excluding Eukaryotes (taxid:2759) as well as the organism that encodes the CRISPR system. All other manipulations of sequence information and prediction of PAM sequences were completed using Spacer2PAM which is available at [INSERT URL TO GITHUB ONCE PUBLIC]. Spacer2PAM requires the following dependencies: dplyr, ggplot2, ggseqlogo(30), taxonomizr, HelpersMG, httr, jsonlite, spatstat.utils, and seqinr. Prophage prediction uses the Phaster API(31). More information about Spacer2PAM can be found in the program documentation.

### Plasmid Construction

All individual plasmids and libraries in this work were generated by two-piece Gibson assembly using the GeneArt Seamless Plus kit. Linear backbone was generated by PCR of pMTL82254 using Kapa DNA polymerase Master Mix and purification by gel electrophoresis and extraction with Zymoclean Gel DNA recovery Kit. Linear dsDNA gBlocks ordered from IDT containing the PAM sequence upstream of *C. autoethanogenum* CRISPR array 1 spacer 19 were used as inserts. Gibson assembly products were transformed into chemically competent One Shot™ MAX Efficiency™ DH10B T1 Phage-Resistant Cells using standard procedures. DNA sequence was confirmed by Illumina MiSeq Sequencing V2 and V3 chemistry.

### Spacer2PAM-informed PAM Prediction Screening

Spacer2PAM was applied to the type I-B CRISPR system of C. autoethanogenum using the comprehensive method. The top 25% of high scoring PAM predictions were used to determine a set of 16 four nucleotide PAM sequences that are likely to be functional. The Spacer2PAM-informed, unpooled PAM library constructs were transformed into *E. coli* HB101 carrying R702(32) (CA434(33)) in parallel. Conjugation of library members into *C. autoethanogenum* DSM 19630, a derivate of type strain DSM 10061, was performed as described earlier(33, 34) using erythromycin (250 µg/mL) and clarithromycin (5 µg/mL) for plasmid selection in *E. coli* and *C. autoethanogenum*, respectively, and trimethoprim (10 µg/mL) as counter selection against *E. coli* CA434. Optical density of donor *E. coli* cultures were measured prior to addition to *C. autoethanogenum* cells. Transconjugant colonies were counted following 4 days of incubation at 37°C under 1.7 × 10^5^ Pa gas (55% CO, 10% N2, 30% CO2, and 5% H2) in gas-tight jars. This was performed in biological triplicate, with 3 separate cultures of donor *E. coli* conjugated to aliquots of a single *C. autoethanogenum* culture.

### Randomized PAM Library Screening

The randomized, pooled PAM library was transformed into NEBExpress® *E. coli* and then purified by QIAprep Spin Miniprep Kit. An aliquot of this DNA was saved to determine PAM frequencies before exposure to the CRISPR system. Electroporation into *C. autoethanogenum* was performed as described previously(35, 36). Following recovery, cells were pelleted by centrifugation at 4000 X g for 10 minutes, 9.5 mL of supernatant was discarded, and cells were resuspended in 500 µL YTF. Resuspensions were split by volume and spread on YTF 1.5% agar supplemented with 5 µG/mL clarithromycin, allowed to dry for ∼30 minutes, and incubated at 37°C for 4 days under 1.7 × 10^5^ Pa gas (55% CO, 10% N2, 30% CO2, and 5% H2) in gas-tight jars. 2.5 mL of Luria broth was added to each plate and plates were scraped. Total DNA from the cell suspension was purified using the MasterPure™ Gram Positive DNA Purification Kit. PCR across the PAM and spacer was performed using Kapa DNA polymerase Master Mix followed by purification by gel electrophoresis (1.5% agarose) and extraction with Zymoclean Gel DNA recovery Kit. Extracts were quantified by Quant-iT (Thermo Fisher Scientific), diluted to 1 ng/uL, and prepared for sequencing following the Illumina 16S amplicon protocol starting at the Index PCR step https://support.illumina.com/documents/documentation/chemistry_documentation/16s/16s-metagenomic-library-prep-guide-15044223-b.pdf. Ampure XP purified libraries were quantified by Quant-iT and sequenced using MiSeq Reagent Kit V3. Frequency of each PAM was determined by counting the occurrence of each PAM next to a correct protospacer sequence within the read. Briefly, all sequence reads are searched for the presence of the C. autoethanogenum Array 1 spacer 19 sequence and are binned as a forward read, reverse read, or does not contain the spacer. For all reads in the forward and reverse bins, the immediate 4 nucleotides upstream or downstream, respectively, are extracted. The sequences extracted from reverse reads are converted to their reverse complement to be compatible with the sequences extracted from forward reads and the two sets of sequences are combined. The frequency of each 4-nucleotide sequence in the combined list is then counted and recorded. The frequency of each PAM was converted to a relative frequency within the total library and the log_2_-fold change in relative frequency was calculated from exposure to the CRISPR system.

## Results

### Spacer2PAM predicts functional PAMs from CRISPR array spacers

We set out to develop a computational framework for predicting functional PAMs from CRISPR array spacers. This framework, which we implement as a comprehensive R package, is called Spacer2PAM (Figure 1). With input of the CRISPR system’s host organism and CRISPR array spacer, Spacer2PAM performs a series of steps of sequence alignment and checks to output a PAM prediction from the alignments (see Supplementary Note 1). At the core of Spacer2PAM is an algorithm, join2PAM, which subjects the aligned sequences to six user set filtering steps to down select the number of alignments that are used in PAM prediction and improve the quality of PAM predictions. The first filter removes redundant alignments and any alignments to the organism that encodes the CRISPR system of interest. Removal of these alignments is important as their presence during prediction will return the CRISPR array repeat as the predicted PAM. The second through fifth filters remove alignments based on the number of gaps present in the alignment, E value of the alignment, the length of the alignment, and the start of the query sequence relative to the spacer sequence start, respectively. The sixth filter is optional, and filters based on whether the alignment occurs in a predicted prophage region in the query genome. Spacer2PAM then outputs a consensus PAM sequences and associated PAM score which is calculated by scaling the number of unique alignments *h*_*unique*_ that were used to generate the consensus PAM prediction by the proportion of possible information content that the consensus PAM encodes as shown by:

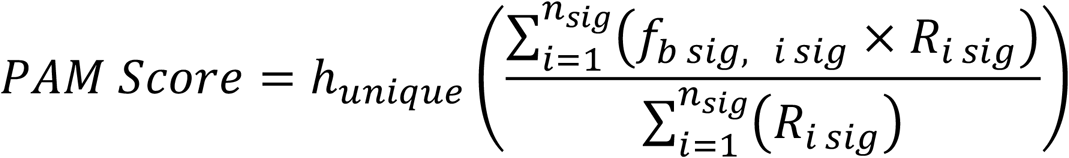

where *n*_*sig*_ is the number of significant nucleotide positions, *f*_*b sig, i sig*_ is the relative frequency of a predicted base *b* at significant position *i*, and *R*_*i sig*_ is the total information content encoded at significant position *i*. For example, if 25 alignments were used to generate a consensus PAM of CC and all 25 alignments encoded the CC motif, the resulting score would be close to 25. If there was disagreement between the sequences in the position of that predicted CC motif, the PAM score would decrease as those two positions would encode less total information content and the C in each position would occur at lower relative frequency. Spacer2PAM can also output a sequence logo of the upstream and downstream PAM predictions using the ggseqlogo package(30) and annotate it with the consensus PAM sequence and PAM score.

**Figure 1.**
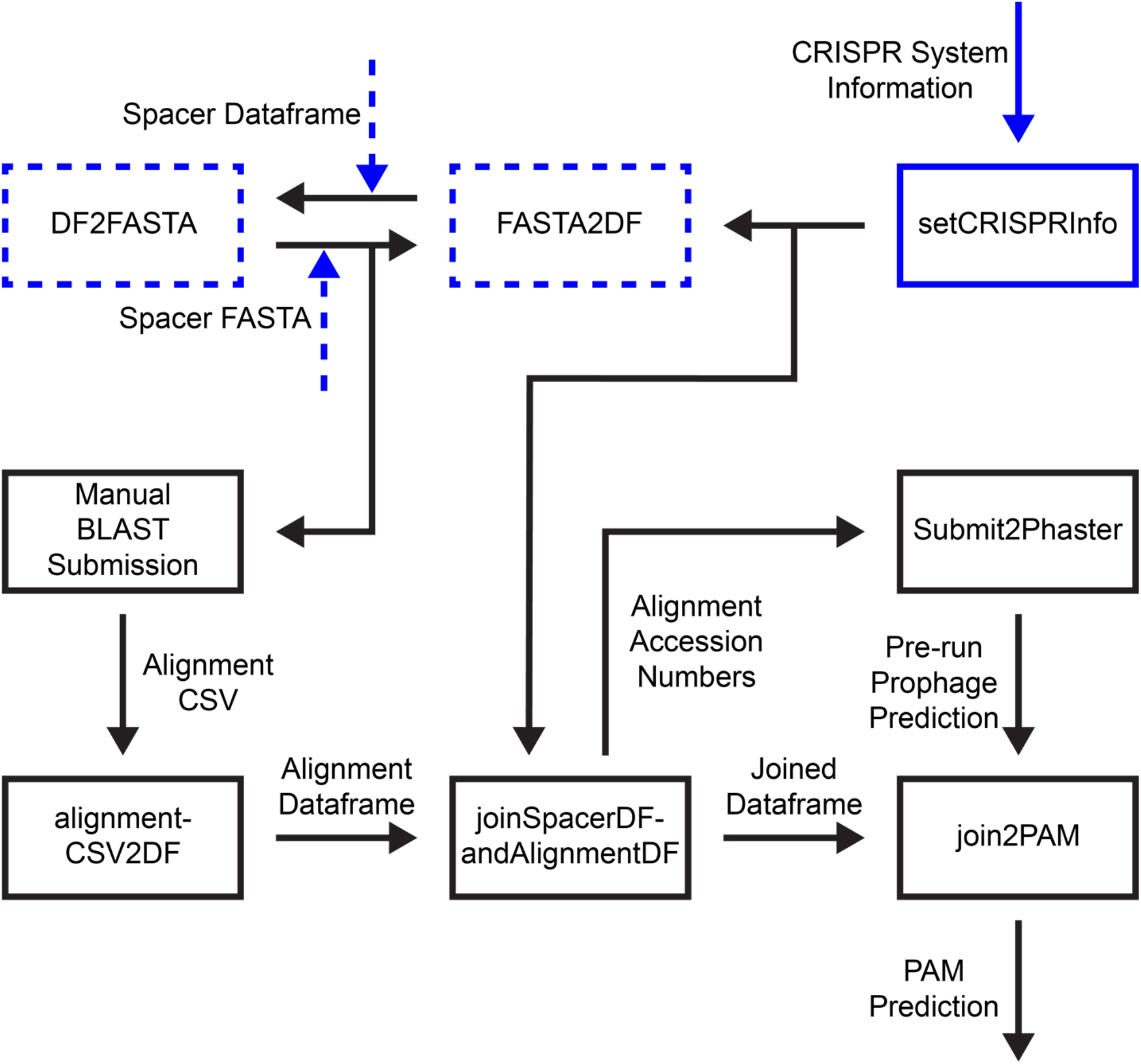
Overview of Spacer2PAM package functions. Functions are represented by boxes and data are represented by arrows. The user should start by inputting information about the CRISPR system via setCRISPR info and by supplying either a FASTA or CSV file containing spacer information. After use of the FASTA file for manual submission to BLAST, functions in Spacer2PAM are used to complete the rest of the data transformations and PAM analysis.

Spacer2PAM was validated by predicting PAMs from the CRISPR array spacers of 10 CRISPR systems with known PAMs over a range of 256 filter criteria sets. Spacer2PAM is effective in predicting PAMs (Figure 2). These model CRISPR effectors have known PAM sequences and come from: *Acinetobacter baumanii*(37), *Bacillus halodurans*(38), *Campylobacter jejuni*(39), *Clostridiodes difficile*(40), *Clostridium pasteurianum*(17), *Clostridium tyrobutyricum*(41), *Hungateiclostridium thermocellum*(16), *Neisseria meningitidis*(24), *Pseudomonas aeruginosa*(42), and *Streptococcus pyogenes*(43). Out of the best PAM predictions for the 10 model systems used, Spacer2PAM predicted functional PAMs for 8. Functional PAMs are defined by sequences that would lead to interference in the presence of the CRISPR system, but the motif may be more restrictive than the true minimal PAM. The best predictions for the remaining 2 model systems yielded partial PAMs, meaning that the prediction is not functional but correctly identifies some positions and residues in the PAM without misidentifying any essential residues. Although these sequences are not functional, they still indicate part of the functional PAM and are valuable in limiting the nucleotide search space. From this analysis, there do not appear to be trends in how well Spacer2PAM performs based on CRISPR system type, however the number of spacers and alignments seem to affect the prediction. Additionally, no incorrect PAM predictions were observed in this sample set.

**Figure 2.**
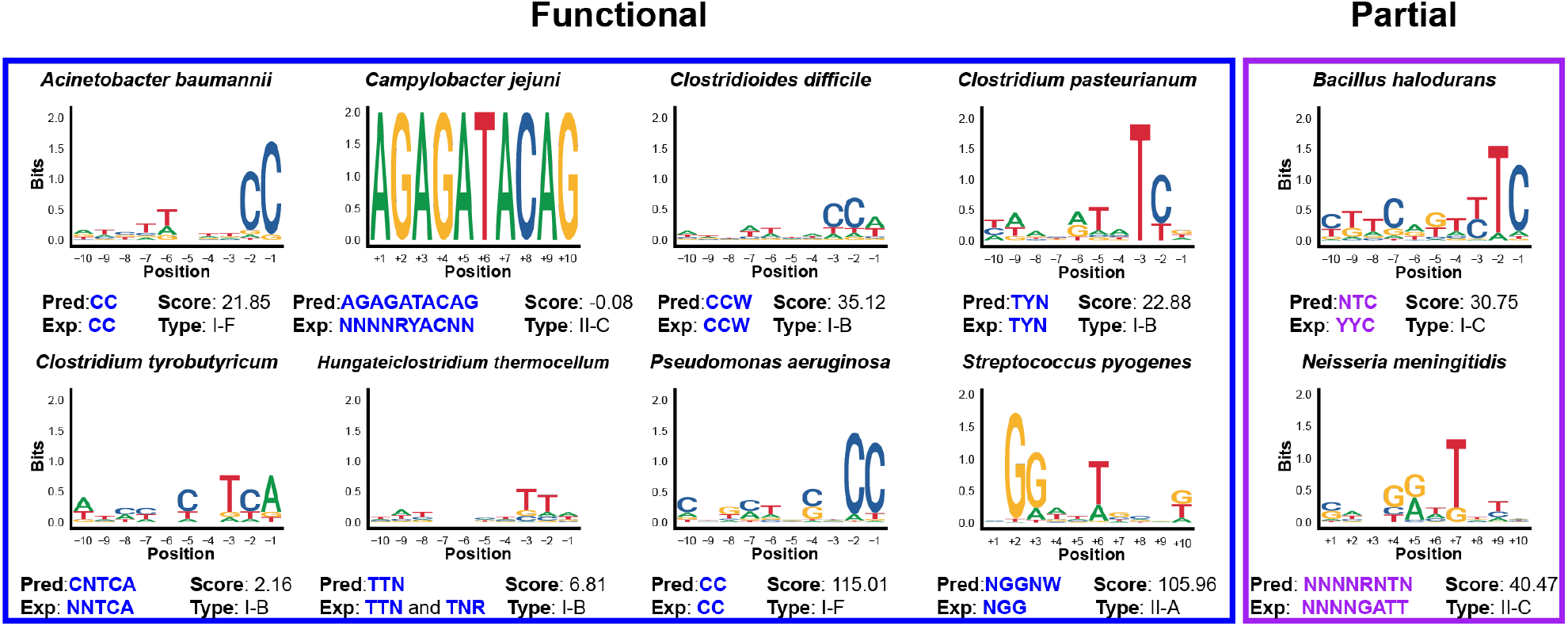
Spacer2PAM recapitulates PAMs from characterized CRISPR systems. Representative sequence logo of the most accurate 10-nucleotide PAM prediction for each of ten CRISPR systems are shown. Predicted sequence, experimentally determined sequence, Spacer2PAM score, and CRISPR system type are indicated for each system. Functional (which are capable of mediating interference) and partial (which do not mediate interference, but do not misidentify any residue) predictions are outlined in blue and purple, respectively.

### Optimization of alignment filter criteria to improve Spacer2PAM performance

Though Spacer2PAM can predict functional PAMs for most of the CRISPR systems evaluated, the filter criteria that yielded the best result in each case varied between organisms. To determine generalized protocols in which Spacer2PAM should be used, we analyzed the outcome of all 256 sets of filter criteria (Figure 3A) for all 10 model CRISPR systems. In doing so, we define two ways in which Spacer2PAM can be used to inform PAM sequences for a given CRISPR system: “Quick” or “Comprehensive.”

**Figure 3.**
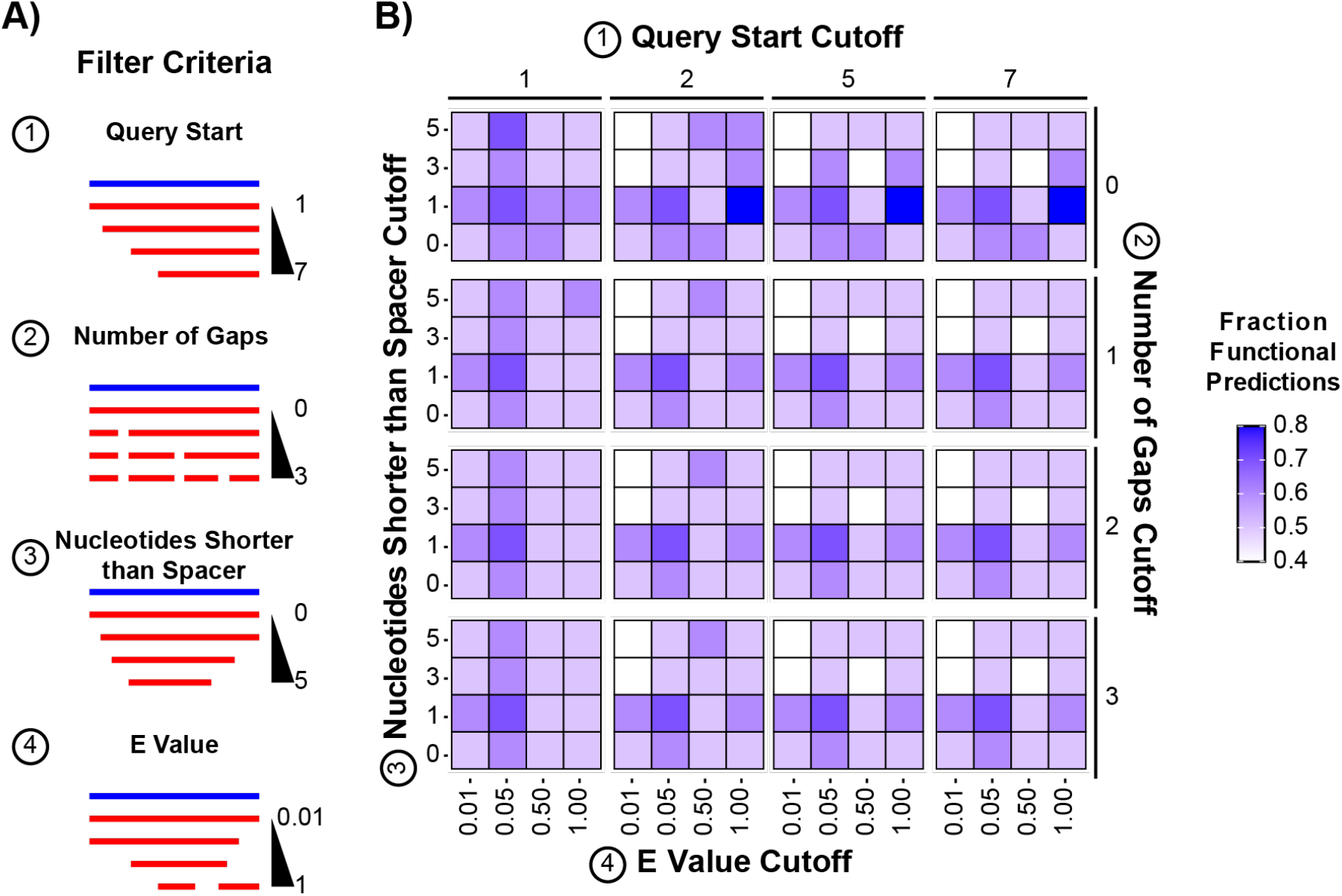
Optimization of filter criteria enables generalized, “Quick” prediction of functional PAMs. Data were generated by filtering alignments to 10 CRISPR systems with known PAMs through 4 filters with 4 different cutoff values. A) Visual representations of each filter criterion. The blue line represents the spacer sequence and the red line represents the query sequence identified by BLAST. The Nucleotides Shorter than Spacer cutoff indicates the threshold value for the difference in alignment and spacer length. The Query Start cutoff indicates the threshold for the starting position of the alignment relative to the spacer. E Value (from BLAST) and Number of Gaps cutoffs are as their names imply. B) The fraction of PAM predictions that resulted in functional sequences out of total predictions is indicated by the fill of each tile.

If computational time or experimental resources are limited, Spacer2PAM can be used in a “Quick” method with optimized filter criteria to suggest a single consensus sequence that is likely to be functional. The filter set chosen for down selecting alignments changes the accuracy of the PAM prediction. With the optimal filter set, Spacer2PAM predicted functional PAMs for 80% of CRISPR systems tested and the remaining 20% of predictions were partial matches for the known PAM (Figure 3B). If predicting a single PAM and not designing a targeted library, the user should use the following filter criteria: E Value cutoff of 1.00, Nucleotides Shorter than Spacer cutoff of 1, Number of Gaps cutoff of 0, and Query Start cutoff of 2. Using a Query Start cutoff of 5 or 7 performs equivalently to a cutoff of 2 in the sample set, but generally a stricter query start cutoff yields better predictions. It is worth noting that using this approach the PAM predicted is more likely to be functional, but also more restrictive than the true minimal PAM consensus.

Alternatively, Spacer2PAM can also be used in a “Comprehensive” method to inform targeted PAM library design if computational time and experimental resources are available. By generating PAM predictions over a range of filter criteria, Spacer2PAM can explore the likely PAM space of a given CRISPR system more thoroughly than single filter set prediction can. Each prediction produces a consensus sequence and is assigned a PAM score which can be used to classify whether an individual PAM prediction should be considered for informing library design. Above a 75^th^ percentile threshold, PAM predictions for the CRISPR systems evaluated were all at least partial matches to the known PAM (Figure 4A). When evaluating the PAM predictions in this scoring bracket, a targeted PAM library can be designed that holds positions supported by multiple predictions constant and varying other positions. This allows the user to change from a pooled, randomized library approach to experimentally simplified unpooled, defined, Spacer2PAM-informed library approach. Additionally, there is often diversity in the PAM prediction using a 75^th^ percentile threshold, allowing for better identification of functional, but divergent PAMs. When this method was applied to the 10 model CRISPR systems, functional PAMs were identified in 90% of the proposed libraries and 70% of the libraries resulted in more than one functional sequence (Figure 4B).

**Figure 4.**
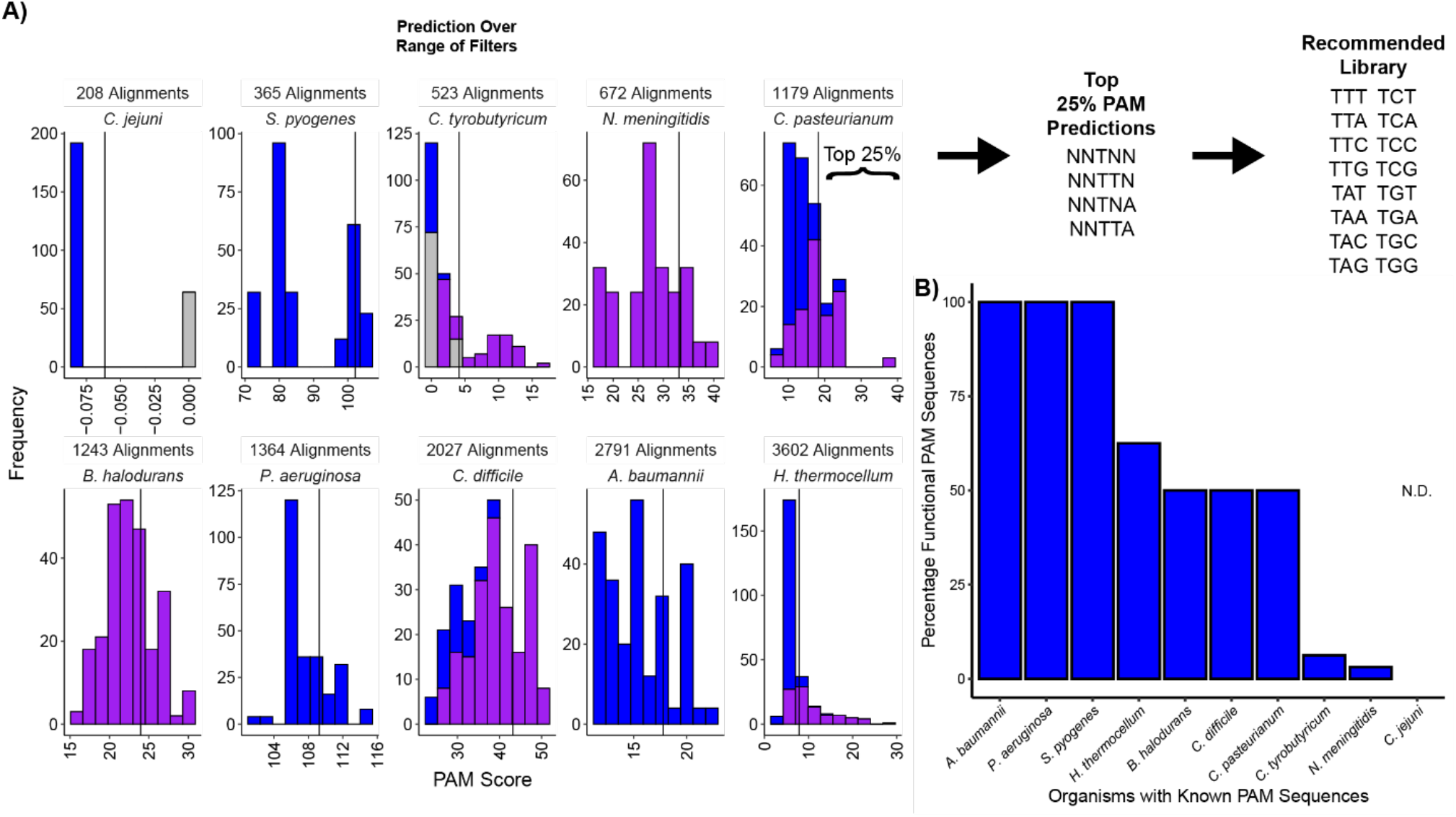
PAM score guides “Comprehensive” PAM prediction. Data were generated by computing PAM predictions and scores over 256 sets of filter criteria for ten CRISPR systems. A) Frequency is plotted against PAM Score for each system. The solid vertical line denotes the 75% percentile PAM score threshold for each CRISPR system. Blue, purple, and gray bars indicate functional, partial, and incorrect PAM predictions, respectively. The top 25% of PAM predictions seed the recommended PAM library for testing. B) Percentage functional PAM sequences within the recommended library are plotted for each CRISPR system determined by comparing known PAM motifs with members of the Spacer2PAM-informed library.

### Application of Spacer2PAM for uncharacterized CRISPR systems

In order to evaluate the efficacy of the generalized protocols for Spacer2PAM, we applied both the “Quick” and “Comprehensive” methods to CRISPR systems with known and unknown PAM sequences. Out of the four characterized CRISPR systems from *Thermobifida fusca* YX, *Clostridium butyricum* JKY6D1, and *Zymomonas mobilis* ZM4 we tested, Spacer2PAM predicted functional PAM sequences for three of them using the “Quick” method and all four with the “Comprehensive” method (Table 1). Both methods were then applied to a variety of uncharacterized CRISPR systems occurring in organisms with unusual carbon metabolism (Table 1). These organisms could be used to convert carbon waste into valuable products. Identifying PAM sequences for their endogenous CRISPR systems could allow for genetic manipulation and genome modification to optimize these organisms for industrial biotechnology.

**Table 1.** Prediction of PAM sequences for organisms with uncommon carbon metabolism. CRISPR spacers and array direction data were downloaded from CRISPRCasdb. Refer to supplementary table S1 for a complete version of this table.

| Organism | CRISPR Type | Quick Prediction | S2P-Informed Recommended Library | Known PAM |
| --- | --- | --- | --- | --- |
| *Thermobifida fusca* YX | III-B | No PAM Predicted | - | No PAM |
|  | I-E | NNNNNNNA/GA/GG | NRRG | WAK |
| *Clostridium butyricum* JKY6D1 | I-B | AANNNNNNCN | TNN | TAA & ACA* |
| *Zymomonas mobilis* ZM4 | I-F | AAGAACTGCC | NCN | CC |
| *Clostridium autoethanogenum* DSM 10061 | I-B | NNNNNNA/TTNA/T | TTNN | N.D. |
| *Clostridium beijerinckii* a4a6934 | I-B | GTTAGCTTTT | NTNT | N.D. |
| *Clostridium saccharoperbutylacetonicum* N1-504 | I-B | ATTTATGTCA | NNCA | N.D. |
| *Methylobacillus flagellatus* KT | I-C | NNNNNNTTTN | NTTN | N.D. |
| *Methylocystis heyeri* H2 | II-C | GCGCCACCGA | ----GNNG | N.D. |
| *Amycolatopsis* sp. BJA-103 | I-E | NGNNNGNNG | NGNG | N.D. |
| *Caldicellulosiruptor bescii* DSM 6725 | Group I Repeat** | NNNNNTNNCA | NNA | N.D. |
|  | Group II Repeat** | NGNNNNNTA | NTN | N.D. |

\*PAM is functional, but not all functional PAMs have been determined
\*\*Arrays were grouped based on the nucleotide identity of the repeat sequence

We further sought to validate the PAM predications from Spacer2PAM for the industrially relevant *Clostridium autoethanogenum. C. autoethanogenum* is an obligate anaerobe with applications in sustainable chemical synthesis(44, 45). We took two approaches to experimentally determine the PAM preference of *C. autoethanogenum’s* type I-B CRISPR system: (i) screening a 16 member, unpooled, Spacer2PAM-informed library and (ii) screening a 256-member, pooled, randomized 4-nucleotide PAM library in the *C. autoethanogenum* host. Both methods involve exposing the PAM library to the active CRISPR system *in vivo* but differ in how the data are collected and evaluated (Figure 5A). Where the pooled library requires the use of NGS before and after screening to measure PAM frequencies, the unpooled method only requires measuring the concentration of donor cells and the number of resulting colonies. Through the unpooled approach, we identified 7 sequences (TTGA, TTGT, TTTA, TTCG, TTCA, TTCT, and TTCC) that resulted in statistically lower (One-tailed Welch’s T-test, p<0.05) conjugation efficiencies than the non-targeting control PAM (AAAT) (Figure 5B). Reduced conjugation efficiency suggests interference by the endogenous CRISPR system. Using the pooled method, we determined that a consensus sequence of NYCN mediates interference and that there is little nucleotide dependence at the -4 position (Figure 5C). By testing the Spacer2PAM predictions, we were able to determine a set of functional PAMs for use in *C. autoethanogenum*.

**Figure 5.**
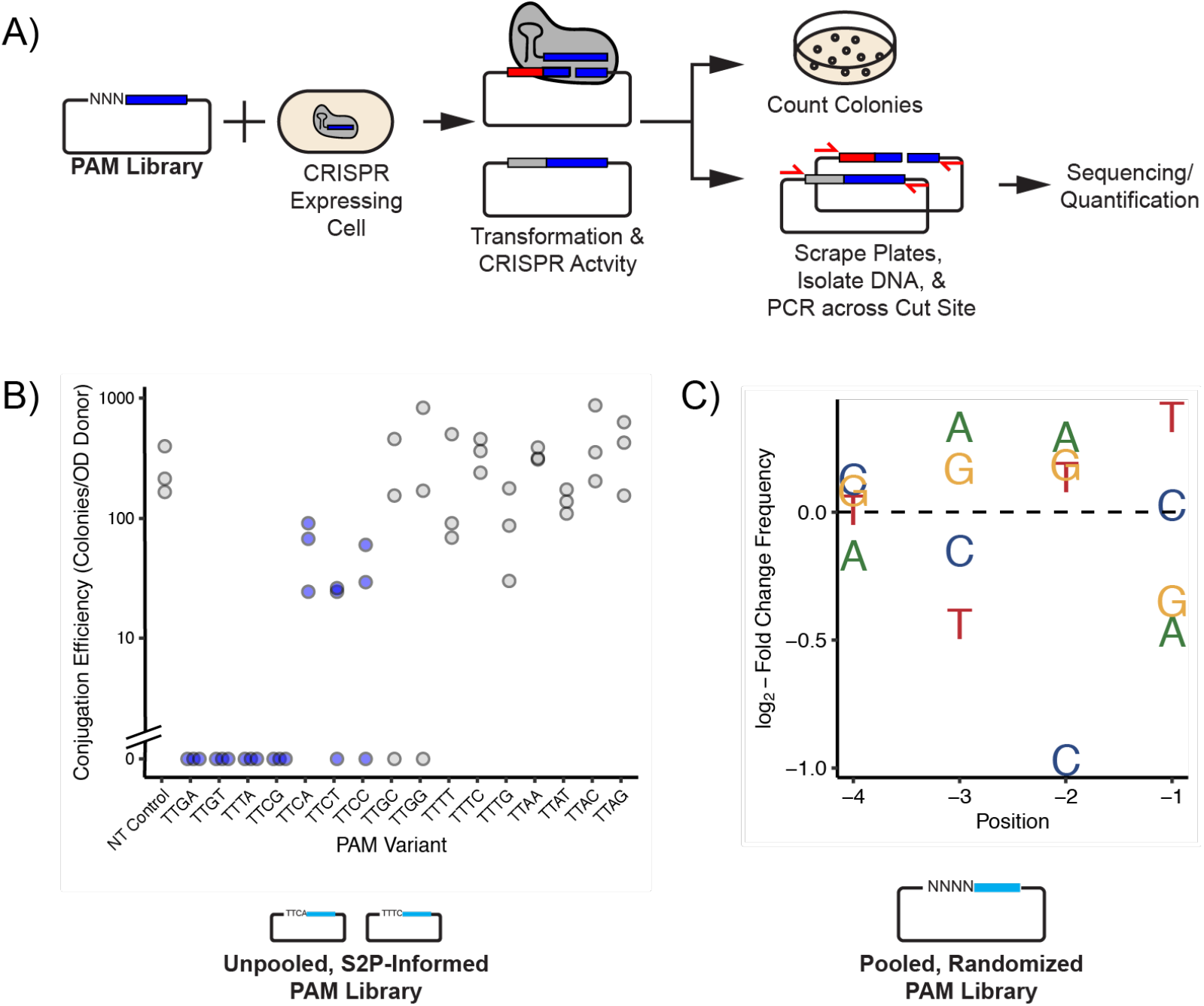
*In vivo* determination of functional PAMs in *C. autoethanogenum*. A) PAM libraries were exposed to active CRISPR systems *in vivo* and then plated on selective media. Readout varied based on library approach. B) An unpooled TTNN PAM library was screened individually by conjugation plasmid from *E. coli* to *C. autoethanogenum*. The non-targeting control PAM was AAAT. Blue indicates p-values less than 0.05 from a one-tailed Welch’s t-test as compared to the non-targeting control. Data are shown in triplicate (n = 3) with three individual experiments, each plotted as a single point. C) A pooled NNNN PAM library was screened *in vivo* by electroporation of plasmid into *C. autoethanogenum*. Nucleotide frequencies were calculated from NGS counts prior and after selection by the CRISPR system.

## Discussion

In this work, we present an easy-to-use, easy-to-interpret computational tool for predicting functional PAM sequences of CRISPR systems. We characterized the tool’s performance to determine 2 methods of use. The “Quick” method uses optimized filter criteria to generate a single consensus PAM using little computational time. The “Comprehensive” method predicts 256 consensus PAMs over a range of filter criteria, which can then be down selected based on PAM score and used to inform a PAM library. The comprehensive method is 90% effective in predicting libraries containing a functional PAM, and both methods narrow the nucleotide search space and allow identification of functional PAMs experimentally more easily. This was exemplified by the ability of a 16-member, Spacer2PAM-informed library to identify 7 functional PAM sequences for the *C. autoethanogenum* type I-B CRISPR system.

Spacer2PAM differs from other computational approaches to PAM prediction in that it employs alignment filtering and produces experimentally actionable outputs. To back track the process of spacer acquisition, Spacer2PAM uses nucleotide alignment through BLAST. While this process is central to the method, nucleotide alignment is inherently sensitive to the length of the sequence submitted. When sequences are short, BLAST is more likely to identify alignments that are not biologically relevant by random chance despite the similarity in nucleotide sequence. As sequences lengthen, the chance of random alignment decreases. Since CRISPR array spacers are relatively short by nature, unfiltered alignments are prone to including biologically irrelevant sequences that then inhibit the ability of PAM prediction programs to identify PAM sequences. Spacer2PAM addresses this by using successive filter criteria to jettison alignments that are less likely to be biologically relevant based on alignments statistics. Though the absolute number of alignments used to generate the consensus PAM decreases, alignments that are likely to lead to a functional PAM are enriched in the filtered subset. Additionally, Spacer2PAM outputs predictions differently than some other programs. While the standard output throughout previous efforts appears to be a sequence logo representative of the potential PAMs used to generate the prediction, interpretation of sequence logos can vary between users. As a result, two researchers may attempt to use divergent PAMs experimentally despite applying the same prediction software. Spacer2PAM still provides the option to generate a sequence logo in addition to the standard output of consensus PAM sequence and PAM score.

In addition to advances in PAM prediction, Spacer2PAM provides a rigorous and reproducible framework in which to choose PAMs for experimental determination. Multiple efforts to functionalize endogenous CRISPR systems for genome engineering have used manual interpretation of BLAST alignments to identify functional PAM sequences(17). Although this approach has yielded success in multiple organisms, it is difficult to reproduce as the researcher makes judgement calls to identify relevant BLAST results. Likewise, the effectiveness of the manual approach is difficult to gauge as it suggested an NAA PAM for the *C. autoethanogenum* type I-B CRISPR system when our work indicates a YCN PAM mediates interference. Using Spacer2PAM and reporting the filter criteria used provides a reproducible way in which to generate and report PAM predictions.

The determination of functional PAMs for the type I-B CRISPR system in *C. autoethanogenum* removes a large hurdle to the functionalization of the system for endogenous genome modification in the organism. While Cas9-based tools have been demonstrated previously in *C. autoethanogenum*(11, 46) and used to vary the metabolic products it produces, the availability of endogenous tools increases the amount of nucleotide cargo that can be delivered while also modulating the genome. Likewise, it is also possible to design and introduce functional synthetic CRISPR arrays into the organism to endow it with resistance to mobile genetic elements such as bacteriophages which have traditionally plagued ABE fermentation processes(47).

We anticipate that the development of Spacer2PAM will encourage the functionalization of endogenous CRISPR systems for a variety of bacteria and archaea as well as help standardize the field. Likewise, Spacer2PAM also has the possibility of streamlining the process of characterizing novel heterologous CRISPR effectors. In both cases, Spacer2PAM represents a step forward that will enable better development of CRISPR technologies for use in prokaryotes and potential acceleration of applied technologies such as CRISPR-based antimicrobials.

## Supporting information

Spacer2PAM Supplement

## Data Availability

Source code for Spacer2PAM as well as instructions are available via GitHub at https://github.com/grybnicky/Spacer2PAM. Illumina sequencing reads for the 4-nucleotide randomized PAM depletion experiment are available through SRA at https://dataview.ncbi.nlm.nih.gov/object/PRJNA755691?reviewer=942vdgmaju9g64bqqv3pqamefu. Further data available on request from the authors.

## Acknowledgements

We would like to sincerely thank Logan Readnour for her help in library preparation and sequencing. We also thank the following investors in LanzaTech’s technology: BASF, CICC Growth Capital Fund I, CITIC Capital, Indian Oil Company, K1W1, Khosla Ventures, the Malaysian Life Sciences, Capital Fund, L. P., Mitsui, the New Zealand Superannuation Fund, Novo Holdings A/S, Petronas Technology Ventures, Primetals, Qiming Venture Partners, Softbank China, and Suncor.

