## Supplementary material for "Spacer2PAM: A computational framework for identification of functional PAM sequences for endogenous CRISPR systems": Spacer2PAM Supplement

**Supplementary Table S1. Extended Prediction of PAM sequences for organisms with uncommon carbon metabolism.** CRISPR spacers and array direction data were downloaded from CRISPRCasdb.

| Organism | CRISPR Type | Array(s) Used | Assumed Direction | Number of Spacers | Number of Alignments | Assumed PAM Location | Quick Prediction | Comprehensive Prediction | Recommended Library | Know PAM |
|---|---|---|---|---|---|---|---|---|---|---|
| *Thermobifida fusca* YX | III-B | 1 | Forward | 3 | 1087 | Upstream | No PAM Predicted | No PAM Predicted | - | No PAM |
| | | 6 | Reverse | 4 | | | | | | |
| | | 10 | Forward | 2 | | | | | | |
| | | 11 | Forward | 4 | | | | | | |
| | | 15 | Forward | 2 | | | | | | |
| | I-E | 3 | Forward | 66 | 15165 | Upstream | NNNNNNNA/GA/G G | NNNNNNNA/GA/GG<br>NNNC/GNNNNNC/G<br>NNNC/GNNC/GNNC/G | NRRG | WAK |
| | | 4 | Reverse | 18 | | | | | | |
| | | 5 | Reverse | 20 | | | | | | |
| | | 7 | Reverse | 27 | | | | | | |
| | | 8 | Reverse | 29 | | | | | | |
| | | 9 | Forward | 11 | | | | | | |
| | | 12 | Reverse | 11 | | | | | | |
| | | 13 | Reverse | 4 | | | | | | |
| | | 16 | Reverse | 13 | | | | | | |
| | | 17 | Reverse | 42 | | | | | | |
| *Clostridium butyricum* JKY6D1 | I-B | 1 | Reverse | 58 | 37 | Upstream | AANNNNNNNCN | NNNNNNNNTNA<br>A/TA/TNNNNNNTNA/C<br>NA/TNNNNA/TTNN<br>NA/TTNNNATNN<br>NA/TNANNA/TA/TNN<br>NA/TTNNNANNN<br>NA/TNA/TNNA/TA/TNN | TNN | TAA & ACA* |
| *Zymomonas mobilis* ZM4 | I-F | 1 | Forward | 8 | 2 | Upstream | AAGAACTGCC | NNGNNNNGCC | NCN | CC |
| | | 2 | Forward | 6 | | | | | | |
| *Clostridium autoethanogenum* DSM 10061 | I-B | 2 | Reverse | 21 | 79 | Upstream | NNNNNNA/TTNA/T | NNNNNNA/TTNN<br>NNNNNNNTNA/T<br>NNNNNNNTNN<br>NNNNNNTTNA/T<br>NNNNNNTTNN<br>NNNTNNTTNN<br>NNTNNNNTNA/T | TTNN | N.D. |
| | | 3 | Reverse | 42 | | | | | | |
| | | 4 | Reverse | 33 | | | | | | |
| *Clostridium beijerinckii* a4a6934 | I-B | 8 | Reverse | 42 | 7 | Upstream | GTTAGCTTTT | NTTNNNNTNT | NTNT | N.D. |
| *Clostridium saccharoperbutylacetonicum* N1-504 | I-B | 8 | Reverse | 27 | 4 | Upstream | ATTTATGTCA | NNNNNNNNCA | NNCA | N.D. |

| | | | | | | | | | | |
|---|---|---|---|---|---|---|---|---|---|---|
| *Methylobacillus flagellatus* KT | I-C | 2 | Forward | 92 | 26 | Upstream | NNNNNNTTTN | A/TT/GNNNNA/TNTT/C<br>NNNNNNNNTC<br>NNNNNNNTTC<br>NNNNNNNTTN<br>NNNNNNTTNN<br>NNNNNNTTTN | NTTN | N.D. |
| *Methylocystis heyeri* H2 | II-C | 3 | Reverse | 11 | 720 | Downstream | GCGCCACCGA | CGAGGGCATT<br>GCGCCACCGA<br>NC/GNNGNNGNN<br>NCC/GNNNNC/GNN<br>NCNNNNNNNN<br>NNNNNNNNNN<br>NNNNNNNNNT | ----GNNG | N.D. |
| *Amycolatopsis sp.* BJA-103 | I-E | 1 | Reverse | 6 | 1380 | Upstream | NGNNNNGNNG | NT/GNA/CA/GNNNNC/G<br>NNNNNNGNNG<br>C/GT/GNNA/GNC/GA/GNC/G<br>NT/GNNA/GNNGNC/G | NGNG | N.D. |
| | | 2 | Reverse | 12 | | | | | | |
| | | 3 | Reverse | 9 | | | | | | |
| *Caldicellulosiruptor bescii* DSM 6725 | Group I Repeat** | 1 | Reverse | 133 | 8139 | Upstream | NNNNNTNNCA | NNNT/GNA/TNNNN<br>NNNNNTNNNA<br>NNNA/TNTNNNN | NNA | N.D. |
| | | 2 | Reverse | 19 | | | | | | |
| | Group II Repeat** | 3 | Forward | 2 | 7468 | Upstream | NGNNNNNNTA | NNNNNNNNTN | NTN | N.D. |
| | | 4 | Forward | 52 | | | | | | |
| | | 5 | Forward | 39 | | | | | | |
| | | 6 | Forward | 2 | | | | | | |
| | | 7 | Forward | 12 | | | | | | |
| | | 8 | Forward | 35 | | | | | | |
| | | 9 | Forward | 5 | | | | | | |

**Supplementary Table S2. Frequencies of 4-nucleotide PAMs before and after selection by the *C. autoethanogenum* type I-B CRISPR system.** Frequencies are normalized to the total number of sequencing reads in each experiment.

| PAM | Fraction of Initial Library | Fraction of Selected Library | log2 Fold Change |
|---|---|---|---|
| AAAA | 0.00314129 | 0.00604512 | 0.94441509 |
| AAAC | 0.0021624 | 0.00130395 | -0.7297451 |
| AAAG | 0.01005801 | 0.01020709 | 0.02122671 |
| AAAT | 0.00071896 | 0.00198155 | 1.46265484 |
| AACA | 0.00489812 | 0.00125987 | -1.9589541 |
| AACC | 1.66E-05 | 1.44E-05 | -0.2092109 |
| AACG | 0.00152087 | 0.00079652 | -0.9331203 |
| AACT | 0.01128761 | 0.01474939 | 0.3859147 |
| AAGA | 0.00272097 | 0.00320247 | 0.23506096 |
| AAGC | 0.00375701 | 0.0048242 | 0.3607065 |
| AAGG | 0.00018803 | 8.10E-05 | -1.2152854 |
| AAGT | 0.01368598 | 0.03361975 | 1.29661048 |
| AATA | 0.01458928 | 0.01637215 | 0.16633491 |
| AATC | 0.00091621 | 0.0017714 | 0.95114469 |
| AATG | 4.24E-05 | 4.51E-05 | 0.08922889 |
| AATT | 0.01594608 | 0.02432296 | 0.60911689 |
| ACAA | 0.0005512 | 0.0004777 | -0.2064563 |
| ACAC | 0.01100003 | 0.00394978 | -1.4776644 |
| ACAG | 0.00050143 | 0.00023578 | -1.0886136 |
| ACAT | 0.00153562 | 0.00146387 | -0.0690332 |
| ACCA | 3.13E-05 | 1.23E-05 | -1.3491411 |
| ACCC | 0.00649826 | 0.00216197 | -1.5877049 |
| ACCG | 0.00235228 | 0.00120041 | -0.970528 |
| ACCT | 0.00716375 | 0.0019959 | -1.8436738 |
| ACGA | 0.00467137 | 0.00174783 | -1.4182856 |
| ACGC | 0.00349892 | 0.0075951 | 1.11815807 |
| ACGG | 1.29E-05 | 1.23E-05 | -0.0690332 |
| ACGT | 0.00115033 | 0.00133778 | 0.2177911 |
| ACTA | 0.00436536 | 0.0014895 | -1.5512752 |
| ACTC | 8.30E-05 | 7.79E-05 | -0.0905664 |
| ACTG | 0.00911784 | 0.00357868 | -1.3492627 |
| ACTT | 0.00856295 | 0.01378783 | 0.6872149 |
| AGAA | 0.00173471 | 0.00364224 | 1.07013029 |
| AGAC | 0.00041847 | 0.00010559 | -1.9866887 |

| | | | |
|---|---|---|---|
| AGAG | 2.03E-05 | 1.44E-05 | -0.4987175 |
| AGAT | 0.01000824 | 0.01060381 | 0.08339502 |
| AGCA | 0.00308045 | 0.00104254 | -1.5630328 |
| AGCC | 1.11E-05 | 1.33E-05 | 0.26883645 |
| AGCG | 0.00343993 | 0.00118401 | -1.5386969 |
| AGCT | 0.00369986 | 0.0048324 | 0.38527037 |
| AGGA | 0.00600789 | 0.00933267 | 0.63543043 |
| AGGC | 3.87E-05 | 2.56E-05 | -0.595102 |
| AGGG | 2.58E-05 | 8.20E-06 | -1.6539957 |
| AGGT | 0.00398007 | 0.00337879 | -0.2362878 |
| AGTA | 0.01164156 | 0.0091461 | -0.3480564 |
| AGTC | 0.00109503 | 0.00151922 | 0.47236984 |
| AGTG | 0.00308414 | 0.00134905 | -1.1929187 |
| AGTT | 0.00641346 | 0.00975502 | 0.60504182 |
| ATAA | 0.00160936 | 0.00178678 | 0.15087824 |
| ATAC | 7.93E-05 | 7.59E-05 | -0.0634522 |
| ATAG | 0.00294588 | 0.00227884 | -0.3704002 |
| ATAT | 0.00487784 | 0.00176218 | -1.4688843 |
| ATCA | 0.00742 | 0.0022204 | -1.7405963 |
| ATCC | 4.06E-05 | 1.85E-05 | -1.1361474 |
| ATCG | 0.00147847 | 0.00058432 | -1.3392811 |
| ATCT | 0.01649359 | 0.00528346 | -1.6423523 |
| ATGA | 0.00496817 | 0.00398463 | -0.3182689 |
| ATGC | 0.00880076 | 0.00331523 | -1.4085197 |
| ATGG | 0.00562261 | 0.0017673 | -1.6696902 |
| ATGT | 0.00422341 | 0.00531113 | 0.33061235 |
| ATTA | 0.01354034 | 0.00477397 | -1.5040024 |
| ATTC | 0.00506772 | 0.00175398 | -1.5307079 |
| ATTG | 0.01110695 | 0.00449411 | -1.3053542 |
| ATTT | 0.00063047 | 0.00108867 | 0.78807476 |
| CAAA | 0.00051433 | 0.000653 | 0.34438748 |
| CAAC | 0.0016665 | 0.00089493 | -0.8969819 |
| CAAG | 0.01133923 | 0.01940547 | 0.77514076 |
| CAAT | 0.0049313 | 0.00355921 | -0.4704127 |
| CACA | 0.01046911 | 0.00530703 | -0.9801614 |
| CACC | 4.98E-05 | 4.41E-05 | -0.1752635 |
| CACG | 8.11E-05 | 3.59E-05 | -1.1767894 |
| CACT | 0.00497923 | 0.00922811 | 0.89011115 |
| CAGA | 0.0010305 | 0.00474937 | 2.20438572 |

| | | | |
|---|---|---|---|
| CAGC | 0.00378466 | 0.00369862 | -0.0331751 |
| CAGG | 4.24E-05 | 3.49E-05 | -0.2827399 |
| CAGT | 5.16E-05 | 0.00010559 | 1.03250483 |
| CATA | 0.00120748 | 0.00474322 | 1.97386952 |
| CATC | 0.00688908 | 0.01553976 | 1.17358134 |
| CATG | 9.95E-05 | 0.00010559 | 0.08497225 |
| CATT | 0.00718035 | 0.02019071 | 1.49156666 |
| CCAA | 0.00145819 | 0.00327833 | 1.16877957 |
| CCAC | 2.58E-05 | 4.31E-05 | 0.73832173 |
| CCAG | 2.58E-05 | 3.18E-05 | 0.30020062 |
| CCAT | 0.0030104 | 0.00725988 | 1.26999128 |
| CCCA | 0.00898511 | 0.00257509 | -1.8029106 |
| CCCC | 0.00038713 | 0.00022143 | -0.8059988 |
| CCCG | 0.00103788 | 0.00056997 | -0.8646908 |
| CCCT | 9.03E-05 | 4.31E-05 | -1.0690332 |
| CCGA | 2.03E-05 | 1.44E-05 | -0.4987175 |
| CCGC | 2.03E-05 | 2.26E-05 | 0.15335923 |
| CCGG | 0.00171812 | 0.00203896 | 0.24700064 |
| CCGT | 0.00460316 | 0.00132035 | -1.801704 |
| CCTA | 0.00484466 | 0.0017386 | -1.4784699 |
| CCTC | 0.00089962 | 0.0010651 | 0.24360183 |
| CCTG | 0.00800069 | 0.00549873 | -0.5410266 |
| CCTT | 0.00704393 | 0.00622041 | -0.1793699 |
| CGAA | 3.32E-05 | 2.97E-05 | -0.1585848 |
| CGAC | 0.01058525 | 0.0107299 | 0.019582 |
| CGAG | 0.00172181 | 0.00045003 | -1.9358424 |
| CGAT | 0.00314682 | 0.01532961 | 2.2843557 |
| CGCA | 4.42E-05 | 1.33E-05 | -1.7311636 |
| CGCC | 0.00777026 | 0.00465916 | -0.7378931 |
| CGCG | 0.00235781 | 0.00071553 | -1.7203581 |
| CGCT | 0.00216056 | 0.00187494 | -0.2045583 |
| CGGA | 0.01042671 | 0.01288983 | 0.30594887 |
| CGGC | 0.00028205 | 0.00015889 | -0.8279042 |
| CGGG | 0.00343809 | 0.0026489 | -0.3762103 |
| CGGT | 0.00441329 | 0.00438135 | -0.010477 |
| CGTA | 8.85E-05 | 6.05E-05 | -0.5489602 |
| CGTC | 5.35E-05 | 9.23E-05 | 0.78723133 |
| CGTG | 0.00338094 | 0.00267146 | -0.3397973 |
| CGTT | 0.00949575 | 0.01675759 | 0.81946068 |

| | | | |
|---|---|---|---|
| CTAA | 1.47E-05 | 2.15E-05 | 0.54567665 |
| CTAC | 7.93E-05 | 9.74E-05 | 0.29695008 |
| CTAG | 0.00145635 | 0.00380319 | 1.38485386 |
| CTAT | 0.00013457 | 0.00010046 | -0.4217555 |
| CTCA | 0.00027468 | 8.30E-05 | -1.7259593 |
| CTCC | 0.00044059 | 0.00022245 | -0.9859563 |
| CTCG | 2.95E-05 | 1.33E-05 | -1.1462011 |
| CTCT | 0.00614615 | 0.0025833 | -1.2504711 |
| CTGA | 4.42E-05 | 3.28E-05 | -0.4316033 |
| CTGC | 0.00216424 | 0.00063967 | -1.7584552 |
| CTGG | 1.29E-05 | 1.54E-05 | 0.2528949 |
| CTGT | 0.0005641 | 0.0002286 | -1.3031287 |
| CTTA | 0.01785961 | 0.00618658 | -1.5294859 |
| CTTC | 0.00517833 | 0.00214352 | -1.2725043 |
| CTTG | 4.61E-05 | 1.85E-05 | -1.320572 |
| CTTT | 0.00291823 | 0.00153768 | -0.9243395 |
| GAAA | 1.47E-05 | 2.87E-05 | 0.96071415 |
| GAAC | 3.87E-05 | 4.92E-05 | 0.34600431 |
| GAAG | 0.00495711 | 0.00600411 | 0.27645137 |
| GAAT | 0.00759882 | 0.00953052 | 0.32677998 |
| GACA | 0.0002839 | 0.00012916 | -1.1361474 |
| GACC | 2.95E-05 | 2.56E-05 | -0.2027846 |
| GACG | 0.01395513 | 0.00666224 | -1.0667163 |
| GACT | 0.00362428 | 0.00146387 | -1.3079081 |
| GAGA | 0.00102682 | 0.00056586 | -0.8596498 |
| GAGC | 0.00087934 | 0.0002327 | -1.9179377 |
| GAGG | 0.00463819 | 0.00713174 | 0.62069263 |
| GAGT | 0.0002544 | 0.00010866 | -1.2272448 |
| GATA | 0.00554518 | 0.00711431 | 0.35949032 |
| GATC | 0.00873808 | 0.01446851 | 0.72752718 |
| GATG | 0.00189141 | 0.00224706 | 0.2485763 |
| GATT | 0.00025624 | 0.00047668 | 0.89550506 |
| GCAA | 5.90E-05 | 1.95E-05 | -1.5987133 |
| GCAC | 3.87E-05 | 3.79E-05 | -0.0295048 |
| GCAG | 0.00304358 | 0.00398156 | 0.38756231 |
| GCAT | 0.00044428 | 0.00020502 | -1.1156739 |
| GCCA | 2.03E-05 | 9.23E-06 | -1.1361474 |
| GCCC | 0.0085648 | 0.00256689 | -1.7383954 |
| GCCG | 2.03E-05 | 5.13E-06 | -1.9841443 |

| | | | |
|---|---|---|---|
| GCCT | 3.87E-05 | 1.54E-05 | -1.3320676 |
| GCGA | 1.66E-05 | 1.44E-05 | -0.2092109 |
| GCGC | 0.00114849 | 0.00155715 | 0.43917703 |
| GCGG | 3.32E-05 | 3.18E-05 | -0.0623695 |
| GCGT | 0.01105349 | 0.01845929 | 0.73984479 |
| GCTA | 0.0016831 | 0.00049411 | -1.7682225 |
| GCTC | 5.35E-05 | 4.72E-05 | -0.1810598 |
| GCTG | 2.95E-05 | 1.54E-05 | -0.9397502 |
| GCTT | 0.00204073 | 0.00313276 | 0.61834855 |
| GGAA | 0.00578852 | 0.00498822 | -0.214669 |
| GGAC | 7.37E-05 | 4.20E-05 | -0.8110169 |
| GGAG | 0.00259193 | 0.00074629 | -1.796227 |
| GGAT | 0.00297169 | 0.00268581 | -0.1459257 |
| GGCA | 0.00050696 | 0.00016402 | -1.6280005 |
| GGCC | 7.37E-06 | 9.23E-06 | 0.32328423 |
| GGCG | 0.00349523 | 0.00123732 | -1.4981741 |
| GGCT | 5.16E-05 | 5.95E-05 | 0.2039853 |
| GGGA | 0.00014379 | 0.00027576 | 0.93941937 |
| GGGC | 0.00053277 | 0.00013839 | -1.9447509 |
| GGGG | 0.0008351 | 0.00030343 | -1.4605546 |
| GGGT | 2.95E-05 | 4.72E-05 | 0.67692118 |
| GGTA | 0.00476355 | 0.00426551 | -0.1593166 |
| GGTC | 0.00208682 | 0.0021999 | 0.07613535 |
| GGTG | 0.0012167 | 0.00044798 | -1.4414735 |
| GGTT | 0.01479944 | 0.02317278 | 0.64688862 |
| GTAA | 0.00072817 | 0.00027781 | -1.3902006 |
| GTAC | 0.01926065 | 0.01642238 | -0.2299934 |
| GTAG | 0.0014287 | 0.00212199 | 0.57072178 |
| GTAT | 0.01919245 | 0.03107951 | 0.69542516 |
| GTCA | 0.00406302 | 0.00194465 | -1.0630453 |
| GTCC | 0.00323899 | 0.00102102 | -1.6655373 |
| GTCG | 0.00238177 | 0.00093593 | -1.3475601 |
| GTCT | 7.19E-05 | 7.48E-05 | 0.05778157 |
| GTGA | 1.47E-05 | 1.44E-05 | -0.0392859 |
| GTGC | 0.00082956 | 0.00113173 | 0.44810249 |
| GTGG | 0.00074661 | 0.00035264 | -1.0821541 |
| GTGT | 3.50E-05 | 5.74E-05 | 0.71278664 |
| GTTA | 0.00177711 | 0.00056484 | -1.6536216 |
| GTTC | 0.00059544 | 0.00020707 | -1.5238196 |

| | | | |
|---|---|---|---|
| GTTG | 0.0012591 | 0.00040185 | -1.6476727 |
| GTTT | 0.01121203 | 0.01566789 | 0.48276367 |
| TAAA | 0.00461422 | 0.00504563 | 0.12894559 |
| TAAC | 0.00552122 | 0.00562789 | 0.02760937 |
| TAAG | 0.00283896 | 0.00208099 | -0.4480914 |
| TAAT | 0.0007503 | 0.00055561 | -0.4333767 |
| TACA | 0.01614149 | 0.00733677 | -1.1375577 |
| TACC | 0.00569266 | 0.00830653 | 0.54514262 |
| TACG | 6.45E-05 | 3.59E-05 | -0.8466408 |
| TACT | 0.00740341 | 0.00306408 | -1.2727371 |
| TAGA | 0.0002673 | 0.00010456 | -1.3541245 |
| TAGC | 0.00507878 | 0.00156638 | -1.6970485 |
| TAGG | 0.00409252 | 0.00362071 | -0.1767151 |
| TAGT | 0.00038713 | 0.00166684 | 2.10622525 |
| TATA | 0.00068946 | 0.00616096 | 3.15961404 |
| TATC | 0.01469252 | 0.02579811 | 0.81218352 |
| TATG | 0.00794355 | 0.00694415 | -0.1939859 |
| TATT | 0.00487969 | 0.00354793 | -0.45981 |
| TCAA | 0.00015301 | 7.18E-05 | -1.0923972 |
| TCAC | 0.00609085 | 0.0113419 | 0.8969467 |
| TCAG | 0.00164438 | 0.00473604 | 1.52613646 |
| TCAT | 0.00277628 | 0.00681806 | 1.29620873 |
| TCCA | 0.00624386 | 0.00181241 | -1.7845305 |
| TCCC | 0.00155405 | 0.00044593 | -1.801158 |
| TCCG | 2.03E-05 | 1.54E-05 | -0.3991818 |
| TCCT | 0.00637106 | 0.00204818 | -1.6371874 |
| TCGA | 0.00306202 | 0.00217017 | -0.4966716 |
| TCGC | 0.00586226 | 0.00489699 | -0.2595627 |
| TCGG | 0.00362981 | 0.00314506 | -0.2068054 |
| TCGT | 0.00519676 | 0.00551308 | 0.08524551 |
| TCTA | 0.00077426 | 0.000204 | -1.9242617 |
| TCTC | 0.00013089 | 9.23E-05 | -0.5045348 |
| TCTG | 0.00065259 | 0.0004572 | -0.5133464 |
| TCTT | 0.01771582 | 0.0264183 | 0.57649941 |
| TGAA | 0.00505113 | 0.00377756 | -0.419152 |
| TGAC | 0.01140928 | 0.01220402 | 0.09714862 |
| TGAG | 0.00019172 | 0.0004326 | 1.1740187 |
| TGAT | 0.01320667 | 0.02515331 | 0.92948086 |
| TGCA | 0.00303252 | 0.00089083 | -1.7673003 |

| | | | |
|---|---|---|---|
| TGCC | 0.00380862 | 0.00136546 | -1.4798869 |
| TGCG | 0.00372936 | 0.00128242 | -1.5400553 |
| TGCT | 0.01174295 | 0.00620299 | -0.9207606 |
| TGGA | 0.00372014 | 0.00596618 | 0.68145221 |
| TGGC | 0.00133468 | 0.00395695 | 1.56789847 |
| TGGG | 2.40E-05 | 2.56E-05 | 0.0967757 |
| TGGT | 0.00132362 | 0.0010897 | -0.2805549 |
| TGTA | 5.35E-05 | 6.97E-05 | 0.38284107 |
| TGTC | 0.00491102 | 0.01075553 | 1.13098295 |
| TGTG | 0.00191169 | 0.00641108 | 1.74572255 |
| TGTT | 0.00452205 | 0.00284573 | -0.6681804 |
| TTAA | 0.00351182 | 0.00166889 | -1.0733311 |
| TTAC | 0.00785321 | 0.02002874 | 1.35071669 |
| TTAG | 0.01966438 | 0.0128396 | -0.6149847 |
| TTAT | 0.00222323 | 0.00143311 | -0.6335063 |
| TTCA | 0.00061203 | 0.00015889 | -1.9455558 |
| TTCC | 0.00382522 | 0.00098821 | -1.9526471 |
| TTCG | 0.00382337 | 0.00129062 | -1.5667784 |
| TTCT | 0.00376807 | 0.0010487 | -1.8452298 |
| TTGA | 0.01188674 | 0.00578679 | -1.0385189 |
| TTGC | 0.00183242 | 0.00455152 | 1.31260115 |
| TTGG | 0.00076136 | 0.00029421 | -1.3717318 |
| TTGT | 0.00127753 | 0.00436905 | 1.77396256 |
| TTTA | 0.01634612 | 0.00617018 | -1.405563 |
| TTTC | 0.0142464 | 0.00715942 | -0.9926823 |
| TTTG | 0.00289242 | 0.00092158 | -1.6500931 |
| TTTT | 0.02021373 | 0.02771918 | 0.45554857 |

Supplementary Table S3. Summary of DNAs used in this study.

| Name | Description | Sequence (5' to 3') | Origin |
|---|---|---|---|
| pMBTL82254 | Plasmid capable of replication in *C. autoethanogenum* and *E. coli* | https://benchling.com/s/seq-au8bXqSOG6WTrZlSCBCP | DOI: 10.1016/j.mimet.2009.05.004 |
| GAR136 | Forward primer to pMBTL82254 backbone for Gibson assembly | TACAGCGGCCGCGGTC | This Study |
| GAR137 | Reverse primer to pMBTL82254 backbone for Gibson assembly | GAAGAGGCCCGCACCGAT | This Study |
| GARB014 | gBlock containing randomized 4-N PAM upstream of *C. autoethanogenum* array 1 spacer 19 and pMBTL82254 homlogy | CAGGAAACAGCTATGACCGCGGCCGCTGTAT<br>CCATACGCGTCCATGGAAGCAAAGGTGAAGA<br>ACTGTTTACCGGCGTTGTGCCGATTCTGGTG<br>GAACTGGATGGCGATGTGAACGGTCACAAAT<br>TCAGNNNNTGCCAAAACTCGTCAACGCCTAA<br>TGCACTAATGGCCGATCTCGAGGCCTGCAGA<br>CATGCAAGCTTGGCACTGGCCGTCGTTTTAC<br>AACGTCGTGACTGGGAAAACCCTGGCGTTAC<br>CCAACTTAATCGCCTTGCAGCACATCCCCCTT<br>TCGCCAGCTGGCGTAATAGCGAAGAGGCCC<br>GCACCGATCGCCCTTCCCAA | This Study |
| GARB015 | gBlock containing TTTT PAM upstream of *C. autoethanogenum* array 1 spacer 19 and pMBTL82254 homlogy | CAGGAAACAGCTATGACCGCGGCCGCTGTAT<br>CCATACGCGTCCATGGAAGCAAAGGTGAAGA<br>ACTGTTTACCGGCGTTGTGCCGATTCTGGTG<br>GAACTGGATGGCGATGTGAACGGTCACAAAT<br>TCAGTTTTTGCCAAAACTCGTCAACGCCTAAT<br>GCACTAATGGCCGATCTCGAGGCCTGCAGAC<br>ATGCAAGCTTGGCACTGGCCGTCGTTTTACA<br>ACGTCGTGACTGGGAAAACCCTGGCGTTACC<br>CAACTTAATCGCCTTGCAGCACATCCCCCTTT<br>CGCCAGCTGGCGTAATAGCGAAGAGGCCCG<br>CACCGATCGCCCTTCCCAA | This Study |
| GARB016 | gBlock containing TTTA PAM upstream of *C. autoethanogenum* array 1 spacer 19 and pMBTL82254 homlogy | CAGGAAACAGCTATGACCGCGGCCGCTGTAT<br>CCATACGCGTCCATGGAAGCAAAGGTGAAGA<br>ACTGTTTACCGGCGTTGTGCCGATTCTGGTG<br>GAACTGGATGGCGATGTGAACGGTCACAAAT<br>TCAGTTTATGCCAAAACTCGTCAACGCCTAAT<br>GCACTAATGGCCGATCTCGAGGCCTGCAGAC<br>ATGCAAGCTTGGCACTGGCCGTCGTTTTACA<br>ACGTCGTGACTGGGAAAACCCTGGCGTTACC<br>CAACTTAATCGCCTTGCAGCACATCCCCCTTT<br>CGCCAGCTGGCGTAATAGCGAAGAGGCCCG<br>CACCGATCGCCCTTCCCAA | This Study |
| GARB017 | gBlock containing TTTC PAM upstream of *C. autoethanogenum* array 1 spacer 19 and pMBTL82254 homlogy | CAGGAAACAGCTATGACCGCGGCCGCTGTAT<br>CCATACGCGTCCATGGAAGCAAAGGTGAAGA<br>ACTGTTTACCGGCGTTGTGCCGATTCTGGTG<br>GAACTGGATGGCGATGTGAACGGTCACAAAT<br>TCAGTTTCTGCCAAAACTCGTCAACGCCTAAT<br>GCACTAATGGCCGATCTCGAGGCCTGCAGAC<br>ATGCAAGCTTGGCACTGGCCGTCGTTTTACA<br>ACGTCGTGACTGGGAAAACCCTGGCGTTACC<br>CAACTTAATCGCCTTGCAGCACATCCCCCTTT<br>CGCCAGCTGGCGTAATAGCGAAGAGGCCCG<br>CACCGATCGCCCTTCCCAA | This Study |

| | | | |
|---|---|---|---|
| GARB018 | gBlock containing TTTG PAM upstream of *C. autoethanogenum* array 1 spacer 19 and pMBTL82254 homlogy | CAGGAAACAGCTATGACCGCGGCCGCTGTATCCATACGCGTCCATGGAAGCAAAGGTGAAGAACTGTTTACCGGCGTTGTGCCGATTCTGGTGGAACTGGATGGCGATGTGAACGGTCACAAATTCAGTTTGTGCCAAAACTCGTCAACGCCTAATGCACTAATGGCCGATCTCGAGGCCTGCAGACATGCAAGCTTGGCACTGGCCGTCGTTTTACAACGTCGTGACTGGGAAAACCCTGGCGTTACCCAACTTAATCGCCTTGCAGCACATCCCCCTTTCGCCAGCTGGCGTAATAGCGAAGAGGCCCGCACCGATCGCCCTTCCCAA | This Study |
| GARB019 | gBlock containing TTAT PAM upstream of *C. autoethanogenum* array 1 spacer 19 and pMBTL82254 homlogy | CAGGAAACAGCTATGACCGCGGCCGCTGTATCCATACGCGTCCATGGAAGCAAAGGTGAAGAACTGTTTACCGGCGTTGTGCCGATTCTGGTGGAACTGGATGGCGATGTGAACGGTCACAAATTCAGTTATTGCCAAAACTCGTCAACGCCTAATGCACTAATGGCCGATCTCGAGGCCTGCAGACATGCAAGCTTGGCACTGGCCGTCGTTTTACAACGTCGTGACTGGGAAAACCCTGGCGTTACCCAACTTAATCGCCTTGCAGCACATCCCCCTTTCGCCAGCTGGCGTAATAGCGAAGAGGCCCGCACCGATCGCCCTTCCCAA | This Study |
| GARB020 | gBlock containing TTAA PAM upstream of *C. autoethanogenum* array 1 spacer 19 and pMBTL82254 homlogy | CAGGAAACAGCTATGACCGCGGCCGCTGTATCCATACGCGTCCATGGAAGCAAAGGTGAAGAACTGTTTACCGGCGTTGTGCCGATTCTGGTGGAACTGGATGGCGATGTGAACGGTCACAAATTCAGTTAATGCCAAAACTCGTCAACGCCTAATGCACTAATGGCCGATCTCGAGGCCTGCAGACATGCAAGCTTGGCACTGGCCGTCGTTTTACAACGTCGTGACTGGGAAAACCCTGGCGTTACCCAACTTAATCGCCTTGCAGCACATCCCCCTTTCGCCAGCTGGCGTAATAGCGAAGAGGCCCGCACCGATCGCCCTTCCCAA | This Study |
| GARB021 | gBlock containing TTAC PAM upstream of *C. autoethanogenum* array 1 spacer 19 and pMBTL82254 homlogy | CAGGAAACAGCTATGACCGCGGCCGCTGTATCCATACGCGTCCATGGAAGCAAAGGTGAAGAACTGTTTACCGGCGTTGTGCCGATTCTGGTGGAACTGGATGGCGATGTGAACGGTCACAAATTCAGTTACTGCCAAAACTCGTCAACGCCTAATGCACTAATGGCCGATCTCGAGGCCTGCAGACATGCAAGCTTGGCACTGGCCGTCGTTTTACAACGTCGTGACTGGGAAAACCCTGGCGTTACCCAACTTAATCGCCTTGCAGCACATCCCCCTTTCGCCAGCTGGCGTAATAGCGAAGAGGCCCGCACCGATCGCCCTTCCCAA | This Study |
| GARB022 | gBlock containing TTAG PAM upstream of *C. autoethanogenum* array 1 spacer 19 and pMBTL82254 homlogy | CAGGAAACAGCTATGACCGCGGCCGCTGTATCCATACGCGTCCATGGAAGCAAAGGTGAAGAACTGTTTACCGGCGTTGTGCCGATTCTGGTGGAACTGGATGGCGATGTGAACGGTCACAAATTCAGTTAGTGCCAAAACTCGTCAACGCCTAATGCACTAATGGCCGATCTCGAGGCCTGCAGACATGCAAGCTTGGCACTGGCCGTCGTTTTACAACGTCGTGACTGGGAAAACCCTGGCGTTACCCAACTTAATCGCCTTGCAGCACATCCCCCTTTCGCCAGCTGGCGTAATAGCGAAGAGGCCCGCACCGATCGCCCTTCCCAA | This Study |

| | | | |
|---|---|---|---|
| GARB023 | gBlock containing TTCT PAM upstream of *C. autoethanogenum* array 1 spacer 19 and pMBTL82254 homlogy | CAGGAAACAGCTATGACCGCGGCCGCTGTATCCATACGCGTCCATGGAAGCAAAGGTGAAGAACTGTTTACCGGCGTTGTGCCGATTCTGGTGGAACTGGATGGCGATGTGAACGGTCACAAATTCAGTTCTTGCCAAAACTCGTCAACGCCTAATGCACTAATGGCCGATCTCGAGGCCTGCAGACATGCAAGCTTGGCACTGGCCGTCGTTTTACAACGTCGTGACTGGGAAAACCCTGGCGTTACCCAACTTAATCGCCTTGCAGCACATCCCCCTTTCGCCAGCTGGCGTAATAGCGAAGAGGCCCGCACCGATCGCCCTTCCCAA | This Study |
| GARB024 | gBlock containing TTCA PAM upstream of *C. autoethanogenum* array 1 spacer 19 and pMBTL82254 homlogy | CAGGAAACAGCTATGACCGCGGCCGCTGTATCCATACGCGTCCATGGAAGCAAAGGTGAAGAACTGTTTACCGGCGTTGTGCCGATTCTGGTGGAACTGGATGGCGATGTGAACGGTCACAAATTCAGTTCATGCCAAAACTCGTCAACGCCTAATGCACTAATGGCCGATCTCGAGGCCTGCAGACATGCAAGCTTGGCACTGGCCGTCGTTTTACAACGTCGTGACTGGGAAAACCCTGGCGTTACCCAACTTAATCGCCTTGCAGCACATCCCCCTTTCGCCAGCTGGCGTAATAGCGAAGAGGCCCGCACCGATCGCCCTTCCCAA | This Study |
| GARB025 | gBlock containing TTCC PAM upstream of *C. autoethanogenum* array 1 spacer 19 and pMBTL82254 homlogy | CAGGAAACAGCTATGACCGCGGCCGCTGTATCCATACGCGTCCATGGAAGCAAAGGTGAAGAACTGTTTACCGGCGTTGTGCCGATTCTGGTGGAACTGGATGGCGATGTGAACGGTCACAAATTCAGTTCCTGCCAAAACTCGTCAACGCCTAATGCACTAATGGCCGATCTCGAGGCCTGCAGACATGCAAGCTTGGCACTGGCCGTCGTTTTACAACGTCGTGACTGGGAAAACCCTGGCGTTACCCAACTTAATCGCCTTGCAGCACATCCCCCTTTCGCCAGCTGGCGTAATAGCGAAGAGGCCCGCACCGATCGCCCTTCCCAA | This Study |
| GARB026 | gBlock containing TTCG PAM upstream of *C. autoethanogenum* array 1 spacer 19 and pMBTL82254 homlogy | CAGGAAACAGCTATGACCGCGGCCGCTGTATCCATACGCGTCCATGGAAGCAAAGGTGAAGAACTGTTTACCGGCGTTGTGCCGATTCTGGTGGAACTGGATGGCGATGTGAACGGTCACAAATTCAGTTCGTGCCAAAACTCGTCAACGCCTAATGCACTAATGGCCGATCTCGAGGCCTGCAGACATGCAAGCTTGGCACTGGCCGTCGTTTTACAACGTCGTGACTGGGAAAACCCTGGCGTTACCCAACTTAATCGCCTTGCAGCACATCCCCCTTTCGCCAGCTGGCGTAATAGCGAAGAGGCCCGCACCGATCGCCCTTCCCAA | This Study |
| GARB027 | gBlock containing TTGT PAM upstream of *C. autoethanogenum* array 1 spacer 19 and pMBTL82254 homlogy | CAGGAAACAGCTATGACCGCGGCCGCTGTATCCATACGCGTCCATGGAAGCAAAGGTGAAGAACTGTTTACCGGCGTTGTGCCGATTCTGGTGGAACTGGATGGCGATGTGAACGGTCACAAATTCAGTTGATGCCAAAACTCGTCAACGCCTAATGCACTAATGGCCGATCTCGAGGCCTGCAGACATGCAAGCTTGGCACTGGCCGTCGTTTTACAACGTCGTGACTGGGAAAACCCTGGCGTTACCCAACTTAATCGCCTTGCAGCACATCCCCCTTTCGCCAGCTGGCGTAATAGCGAAGAGGCCCGCACCGATCGCCCTTCCCAA | This Study |

| | | | |
|---|---|---|---|
| GARB028 | gBlock containing TTGA PAM upstream of *C. autoethanogenum* array 1 spacer 19 and pMBTL82254 homlogy | CAGGAAACAGCTATGACCGCGGCCGCTGTAT<br>CCATACGCGTCCATGGAAGCAAAGGTGAAGA<br>ACTGTTTACCGGCGTTGTGCCGATTCTGGTG<br>GAACTGGATGGCGATGTGAACGGTCACAAAT<br>TCAGTTGTTGCCAAAACTCGTCAACGCCTAAT<br>GCACTAATGGCCGATCTCGAGGCCTGCAGAC<br>ATGCAAGCTTGGCACTGGCCGTCGTTTTACA<br>ACGTCGTGACTGGGAAAACCCTGGCGTTACC<br>CAACTTAATCGCCTTGCAGCACATCCCCCTTT<br>CGCCAGCTGGCGTAATAGCGAAGAGGCCCG<br>CACCGATCGCCCTTCCCAA | This Study |
| GARB029 | gBlock containing TTGC PAM upstream of *C. autoethanogenum* array 1 spacer 19 and pMBTL82254 homlogy | CAGGAAACAGCTATGACCGCGGCCGCTGTAT<br>CCATACGCGTCCATGGAAGCAAAGGTGAAGA<br>ACTGTTTACCGGCGTTGTGCCGATTCTGGTG<br>GAACTGGATGGCGATGTGAACGGTCACAAAT<br>TCAGTTGCTGCCAAAACTCGTCAACGCCTAAT<br>GCACTAATGGCCGATCTCGAGGCCTGCAGAC<br>ATGCAAGCTTGGCACTGGCCGTCGTTTTACA<br>ACGTCGTGACTGGGAAAACCCTGGCGTTACC<br>CAACTTAATCGCCTTGCAGCACATCCCCCTTT<br>CGCCAGCTGGCGTAATAGCGAAGAGGCCCG<br>CACCGATCGCCCTTCCCAA | This Study |
| GARB030 | gBlock containing TTGG PAM upstream of *C. autoethanogenum* array 1 spacer 19 and pMBTL82254 homlogy | CAGGAAACAGCTATGACCGCGGCCGCTGTAT<br>CCATACGCGTCCATGGAAGCAAAGGTGAAGA<br>ACTGTTTACCGGCGTTGTGCCGATTCTGGTG<br>GAACTGGATGGCGATGTGAACGGTCACAAAT<br>TCAGTTGGTGCCAAAACTCGTCAACGCCTAA<br>TGCACTAATGGCCGATCTCGAGGCCTGCAGA<br>CATGCAAGCTTGGCACTGGCCGTCGTTTTAC<br>AACGTCGTGACTGGGAAAACCCTGGCGTTAC<br>CCAACTTAATCGCCTTGCAGCACATCCCCCTT<br>TCGCCAGCTGGCGTAATAGCGAAGAGGCCC<br>GCACCGATCGCCCTTCCCAA | This Study |
| GARB031 | gBlock containing AAAT PAM (non-targeting) upstream of *C. autoethanogenum* array 1 spacer 19 and pMBTL82254 homlogy | CAGGAAACAGCTATGACCGCGGCCGCTGTAT<br>CCATACGCGTCCATGGAAGCAAAGGTGAAGA<br>ACTGTTTACCGGCGTTGTGCCGATTCTGGTG<br>GAACTGGATGGCGATGTGAACGGTCACAAAT<br>TCAGAAATTGCCAAAACTCGTCAACGCCTAAT<br>GCACTAATGGCCGATCTCGAGGCCTGCAGAC<br>ATGCAAGCTTGGCACTGGCCGTCGTTTTACA<br>ACGTCGTGACTGGGAAAACCCTGGCGTTACC<br>CAACTTAATCGCCTTGCAGCACATCCCCCTTT<br>CGCCAGCTGGCGTAATAGCGAAGAGGCCCG<br>CACCGATCGCCCTTCCCAA | This Study |
| pGAR075 | pMBTL82254 plasmid containing *C. autoethanogenum* array 1 spacer 19 and upstream NNNN PAM | https://benchling.com/s/seq-FBUNvg3duner1AD1vdIq | This Study |
| pGAR076 | pMBTL82254 plasmid containing *C. autoethanogenum* array 1 spacer 19 and upstream TTTT PAM | Refer to pGAR075 | This Study |
| pGAR077 | pMBTL82254 plasmid containing *C. autoethanogenum* array 1 spacer 19 and upstream TTTA PAM | Refer to pGAR075 | This Study |

| | | | |
|---|---|---|---|
| pGAR078 | pMBTL82254 plasmid containing *C. autoethanogenum* array 1 spacer 19 and upstream TTTC PAM | Refer to pGAR075 | This Study |
| pGAR079 | pMBTL82254 plasmid containing *C. autoethanogenum* array 1 spacer 19 and upstream TTTG PAM | Refer to pGAR075 | This Study |
| pGAR080 | pMBTL82254 plasmid containing *C. autoethanogenum* array 1 spacer 19 and upstream TTAT PAM | Refer to pGAR075 | This Study |
| pGAR081 | pMBTL82254 plasmid containing *C. autoethanogenum* array 1 spacer 19 and upstream TTAA PAM | Refer to pGAR075 | This Study |
| pGAR082 | pMBTL82254 plasmid containing *C. autoethanogenum* array 1 spacer 19 and upstream TTAC PAM | Refer to pGAR075 | This Study |
| pGAR083 | pMBTL82254 plasmid containing *C. autoethanogenum* array 1 spacer 19 and upstream TTAG PAM | Refer to pGAR075 | This Study |
| pGAR084 | pMBTL82254 plasmid containing *C. autoethanogenum* array 1 spacer 19 and upstream TTCT PAM | Refer to pGAR075 | This Study |
| pGAR085 | pMBTL82254 plasmid containing *C. autoethanogenum* array 1 spacer 19 and upstream TTCA PAM | Refer to pGAR075 | This Study |
| pGAR086 | pMBTL82254 plasmid containing *C. autoethanogenum* array 1 spacer 19 and upstream TTCC PAM | Refer to pGAR075 | This Study |
| pGAR087 | pMBTL82254 plasmid containing *C. autoethanogenum* array 1 spacer 19 and upstream TTCG PAM | Refer to pGAR075 | This Study |
| pGAR088 | pMBTL82254 plasmid containing *C. autoethanogenum* array 1 spacer 19 and upstream TTGT PAM | Refer to pGAR075 | This Study |
| pGAR089 | pMBTL82254 plasmid containing *C. autoethanogenum* array 1 spacer 19 and upstream TTGA PAM | Refer to pGAR075 | This Study |
| pGAR090 | pMBTL82254 plasmid containing *C. autoethanogenum* array 1 spacer 19 and upstream TTGC PAM | Refer to pGAR075 | This Study |

| | | | |
|---|---|---|---|
| pGAR091 | pMBTL82254 plasmid containing *C. autoethanogenum* array 1 spacer 19 and upstream TTGG PAM | Refer to pGAR075 | This Study |
| pGAR092 | pMBTL82254 plasmid containing *C. autoethanogenum* array 1 spacer 19 and upstream AAAT PAM | Refer to pGAR075 | This Study |
| GAR138 | Illumina Sequencing forward primer for pGAR075 | TCGTCGGCAGCGTCAGATGTGTATAAGAGACAGGACCGCGGCCGCTGTA | This Study |
| GAR139 | Illumina Sequencing reverse primer pGAR075 | GTCTCGTGGGCTCGGAGATGTGTATAAGAGACAGATCGGTGCGGGCCTCTTC | This Study |
